# Alginate-regulating genes are identified in the clinical cystic fibrosis isolate of *Pseudomonas aeruginosa* PA2192

**DOI:** 10.1101/319004

**Authors:** Brett Colbert, Hansi Kumari, Ana Piñon, Lior Frey, Sundar Pandey, Kalai Mathee

**Affiliations:** Department of Biological Sciences, College of Arts and Sciences, Florida International University, Miami, FL, USA; Department of Human and Molecular Genetics, Herbert Wertheim College of Medicine, Miami, FL, USA

**Keywords:** Sigma factor, anti-sigma factor, cystic fibrosis

## Abstract

Cystic fibrosis (CF) is a genetic disorder that leads to a buildup of mucus in the lungs ideal for bacterial colonization. When *Pseudomonas aeruginosa* enters the CF lung, it undergoes a conversion from nonmucoid to mucoid; colonization by a mucoid strain of *P. aeruginosa* greatly increases mortality. The mucoid phenotype is due to the production of alginate. The regulator of alginate production is the AlgT/U sigma factor. The observed phenotypic conversion is due to a mutation in the *mucA* gene coding for an anti-sigma factor, MucA, which sequesters AlgT/U. This mucoid phenotype is unstable when the strains are removed from the lung as they acquire second-site mutations. This *in vitro* reversion phenomenon is utilized to identify novel genes regulating alginate production. Previously, second-site mutations were mapped to *algT/U, algO,* and *mucP*, demonstrating their role in alginate regulation. Most of these studies were performed using a non-CF isolate. It was hypothesized that second site mutations in a clinical strain would be mapped to the same genes. In this study, a clinical, hyper-mucoid *P. aeruginosa* strain PA2192 was used to study the reversion phenomenon. This study found that PA2192 has a novel *mucA* mutation which was named them *mucA180* allele. Twelve colonies were sub-cultured for two weeks without aeration at room temperature in order to obtain nonmucoid ***s****uppressors of* ***a****lginate* ***p****roduction* (*sap*). Only 41 *sap* mutants were stable for more than 48 hours — a reversion frequency of 3.9% as compared to ~90% in laboratory strains showing that PA2192 has a stable mucoid phenotype. This phenotype was restored in 28 of the 41 *sap* mutants when complemented with plasmids harboring *algT/U*. Four of the *sap* mutants are complemented by *algO*. Sequence analyses of the *algT/U* mutants have found no mutations in the coding region or promoter leading to the hypothesis that there is another, as yet unidentified mechanism of alginate regulation in this clinical strain.

## INTRODUCTION

Cystic fibrosis (CF) is the most common cause of death due to genetic disorder (1). CF is an autosomal recessive disorder caused by a mutation in the cystic fibrosis transmembrane conductance regulator (*cftr)* gene (2-4). The most common CF mutation (~90%) is the ΔF508 which is the deletion of three nucleotides leading to the loss of a phenylalanine in the 508 position, and thus, a defective protein (3). A wide range of other mutations are possible that lead to either impaired function or total loss of activity (5).

Ordinarily, CFTR functions as anion transporter (6). When mutated, normal anion flow is restricted (7) and mucus accumulates in the CF lung; resulting pulmonary failure is the foremost killer of CF patients (8, 9) The mucus buildup provides a breeding ground for many pathogenic bacterial species especially *S. aureus*, *H. influenzae*, *Pseudomonas aeruginosa*, and *Burkholderia cenocepacia* (10); the relative population of each species fluctuates over the life of the individual (11). The pathogen that rises to prominence over the life of a CF patient and is the leading cause of mortality is *P. aeruginosa* (11, 12).

*P. aeruginosa* expresses a multitude of virulence factors (13). The major contributor to *P. aeruginosa* virulence in patients with CF is its ability to change from the standard, non-mucoid form to the mucoid form (14, 15). Mucoid *P. aeruginosa* is considered highly virulent because patients show poor clinical outcome despite having a heightened immune response (14, 15). The mucoid phenotype is a result of the production of a complex polysaccharide called alginate (16).

Alginate protects *P. aeruginosa* from phagocytosis, antibiotics, oxygen radicals, and the host immune response (17-23) Leid et al., 2005). The importance of alginate in the virulence of *P. aeruginosa* has also been demonstrated in mouse models (24, 25). In mice, an alginate-overproducing strain causes aggressive polymorphonuclear leukocyte (PMN) infiltration-similar to human infection- and causes inefficient pulmonary clearance. A protracted lung infection has the potential to spread to other organs such as the spleen. These properties suggest that alginate is an important virulence factor.

Alginate biosynthesis comes at a high metabolic cost, and thus is tightly regulated (Figure 1) by an intricate system of periplasmic and inner membrane proteins (26-29). The primary regulatory unit of alginate production is a five-gene operon containing *algT/U-mucA-mucB-mucC-mucD* (30). The first gene of this operon, *algT/U*, codes for a sigma factor able to bind to RNA polymerase (RNAP), guiding it to transcribe the genes necessary for alginate production (31, 32).

**Figure 1:**
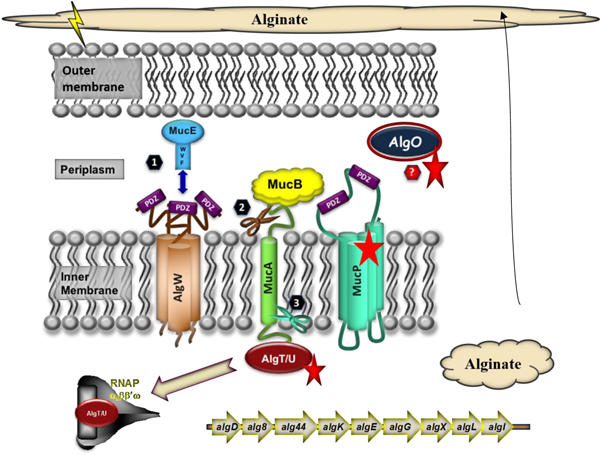
The *Pseudomonas aeruginosa* alginate regulation pathway. Alginate production is controlled by the sigma factor AlgT/U which is ordinarily bound to the inner membrane by the anti-sigma factor MucA to prevent interaction with RNAP. AlgT/U must be freed from MucA to begin alginate production. When stress is sensed MucE misfolds (1) and induces periplasmic cleavage of MucA by AlgW (2). MucA is also cleaved by MucP (3) on the cytoplasmic end to release AlgT/U. AlgT/U is now free to interact with RNAP and initiate alginate biosynthesis by transcribing the *algD* operon. When MucA is mutated, it is unable to sequester AlgT/U and a mucoid phenotype ensues. Proteins marked with a red star are under investigation in this study. Adapted from Pandey *et al.,* 2016 (27).

Under normal circumstances, *P. aeruginosa* is non-mucoid, as is the case with the prototypic reference strain, PAO1 (33). Upon colonizing the CF lung, *P. aeruginosa* must confront the host immune system and antibiotics. The typical response is to convert to a mucoid phenotype by producing alginate. This is commonly accomplished by mutating mucA which codes for the anti-sigma factor to AlgT/U (34). Ordinarily, MucA sequesters AlgT/U to the inner membrane, preventing it from directing RNAP; however, when mucA is mutated, AlgT/U is left free to guide RNAP (Figure 1) to transcribe the genes needed for alginate biosynthesis (27, 35). The most common mucA mutation (~85%) found in clinical, mucoid strains of *P. aeruginosa* is the mucA22 allele which is the deletion of a single G in a string of five Gs resulting in a frameshift mutation and premature stop codon (34, 35).

Since alginate production is metabolically expensive, mucoid strains revert to a non-mucoid phenotype when isolated from the lung and cultured *in vitro* (Figure 2), especially when grown at low oxygen levels (36). The isolates maintain the original *mucA* mutation but revert to Alg^-^ by mutating at another gene, a second site crucial to alginate biosynthesis (32, 37). This has proven to be a highly advantageous phenomenon when it comes to determining novel genes involved in alginate regulation. Several studies have utilized this *in vitro* reversion phenomenon to map second-site mutations to genes coding for the sigma factor AlgT/U (31, 32, 37), a putative periplasmic protease AlgO (37, 38), and an inner membrane protease MucP (Delgado *et al.*, submitted).

**Figure 2:**
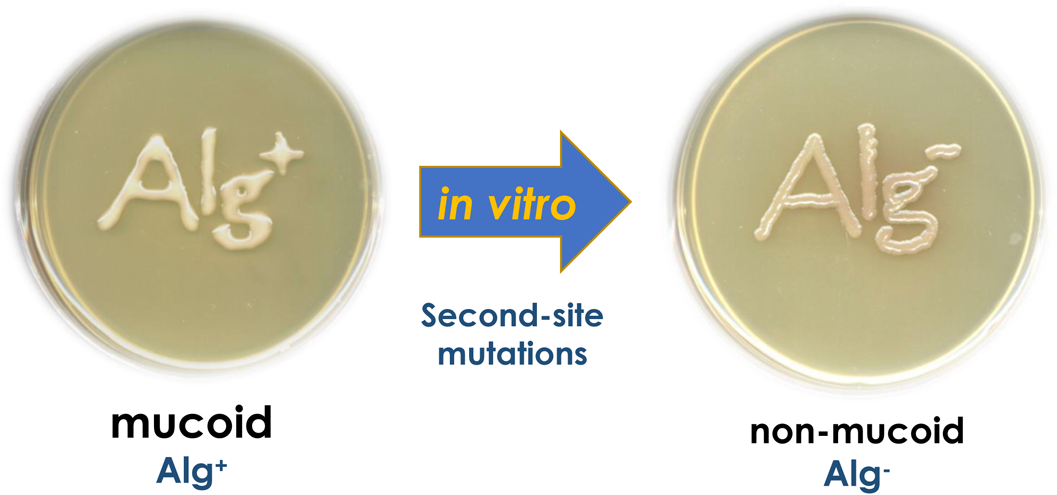
The *P. aeruginosa in vitro* reversion phenomenon. Many clinical isolates are mucoid when removed from the lung due to *mucA* mutations. When cultured *in vitro*, they cease producing alginate and become non-mucoid. *mucA* has been seen to remain mutated, and so the reversion is due to second-site mutations in other alginate-regulating genes.

The study by DeVries et al., (1994) used the mucoid CF isolate FRD1, but the great caveat tied to many of these other studies is that they were carried out in a laboratory-generated strain, PDO300 (22). This form is an isogenic derivative of PAO1 with the addition of the *mucA22* allele in order to imitate clinical isolates (22). The potential issue with using PDO300 is that laboratory strains cannot faithfully mirror the real-world pathogenesis of a clinical isolate (39).

The present study was undertaken to map the location of second-site mutations in a clinical isolate. The strain utilized is *P. aeruginosa* 2192 which was isolated from a CF patient in Boston who passed away from the infection (40). *P. aeruginosa* 2192 produces about 60% more alginate than PDO300 (Delgado *et al.*, submitted) and is far more stable in its mucoid phenotype (this study). We hypothesized that the non-mucoid revertants of *P. aeruginosa* 2192 would harbor second-site mutations in *algT/U*, *algO*, and *mucP* while maintaining the original *mucA* mutation. This would demonstrate the role of these genes in *P. aeruginosa* 2192 alginate regulation.

## MATERIALS AND METHODS

### Bacterial Strains

The *P. aeruginosa* and *Escherichia coli* strains used in this study are listed in Table 1. The *E. coli* strains were grown on Luria-Bertani (LB) media supplemented with tetracycline (Tc) at 20 μg/ml and ampicillin (Ap) at 50 μg/ml. *Pseudomonas aeruginosa* strains were grown on LB or LB/PIA plates, which is a 1:1 mixture of LB and *Pseudomonas* isolation agar. These were supplemented with Tc at 100 μg/ml or carbenicillin (Cb) at 150 μg/ml when appropriate.

**Table 1:**
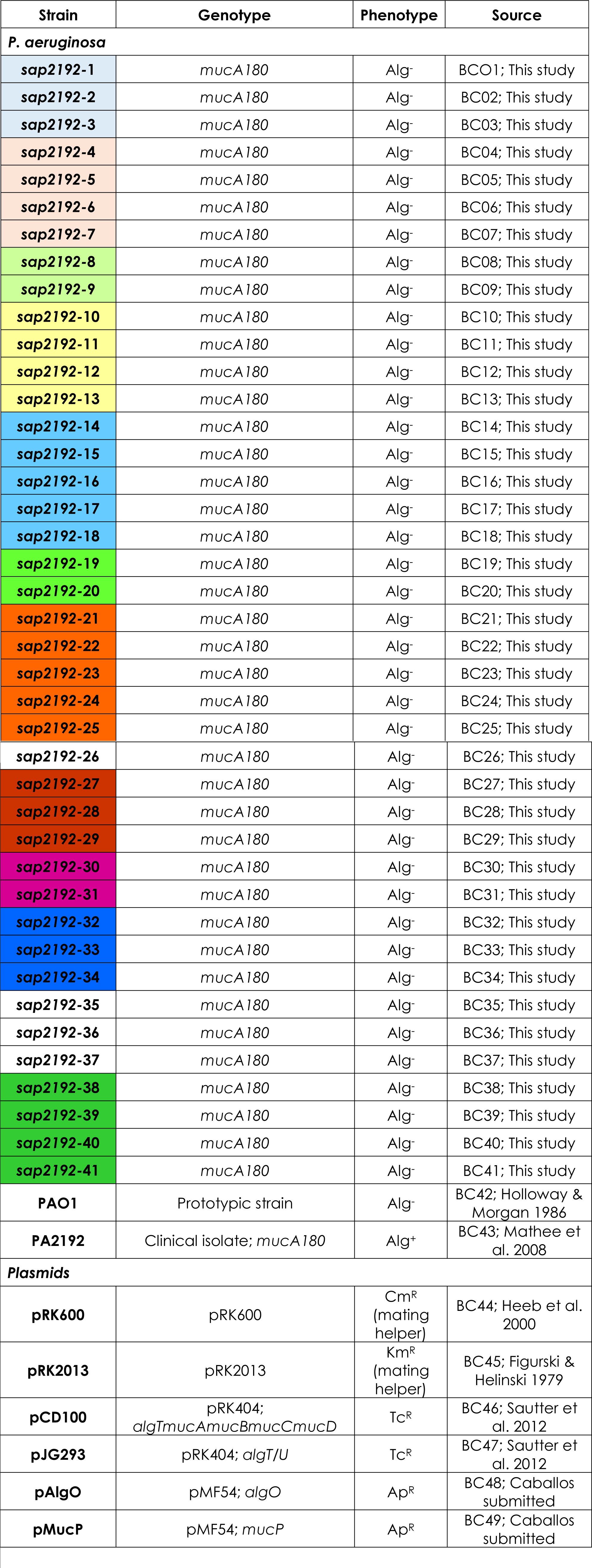
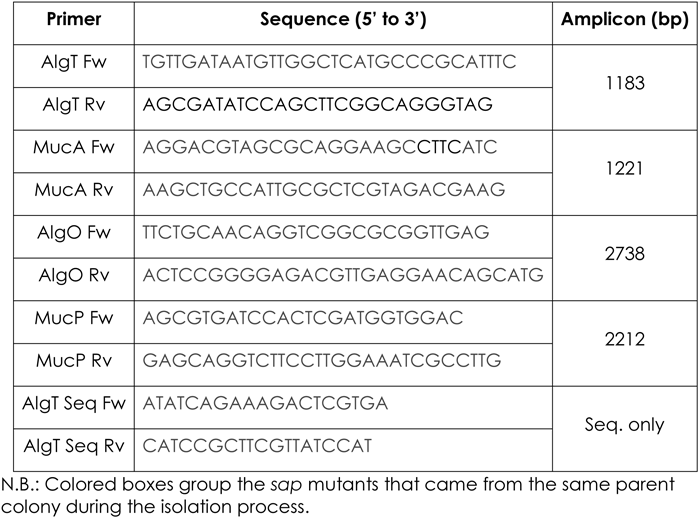
Strains and primers generated and used in this study.

### Isolation of *sap* mutants

The parent strain used to isolate **s**uppressor of **a**lginate **p**roduction (*sap*) mutants was *P. aeruginosa* 2192, a hypermucoid, clinical CF isolate possessing a *mucA* mutation (Mathee *et al.,* 2008, Delgado *et al*., Submitted). *P. aeruginosa* 2192 was plated and grown overnight on LB/agar plates. Twelve mucoid colonies were selected and inoculated into separate tubes containing 5 ml of LB nutrient broth. These were serially cultured at 25°C without aeration for two weeks. Dilutions of each of the 12 cultures were plated daily for single colonies. The 127 *sap* mutant colonies were frozen in 1:1 culture/skim milk at −80°C for further analysis (Table 1). At the end of two weeks, all *sap* mutants were re-streaked on to LB/PIA plates and incubated at 37°C for 24 hours followed by 25°C for 24 more hours. Only the 41 *sap* mutants that maintained a non-mucoid phenotype after 48 hours during this secondary screening were used in subsequent analyses.

### Complementation assays

Complementation of the *sap* mutants was accomplished by a modified tri-parental mating protocol (41) developed during this study. The donor *E. coli* strain containing the plasmid of interest was crossed on an LB plate with the two helper *E. coli* strains, pRK600 and pRK2013 (Table 1), and the recipient, in this case, the *sap* mutants. The following day, the mating conglomeration was homogenized in LB broth, diluted, and plated on selective media (LB/PIA supplemented by Tc 100 μg/ml or Cb 150 μg/ml). Colonies were checked for a mucoid or non-mucoid phenotype at 24 and 48 hours. Each of the *sap* mutants was complemented with pCD100 (Tc resistant) which contains the *algT*/*U-mucA22-mucB-mucC-mucD* operon (37). Those that saw a reversion to a mucoid phenotype were then complemented by pJG293 (Tc resistant) which contains *algT*/*U* alone (37). The remaining *sap* mutants that were not complemented by *algT*/*U* were conjugated with a plasmid containing *algO* (pAlgO), and another harboring *mucP* (pMucP).

### Genomic DNA isolation

Genomic DNA was isolated from each of the *sap* mutants following a standard phenol-chloroform protocol (42). Briefly, 1 ml of an overnight culture was pelleted and mixed with lysozyme and proteinase K. After incubation at 37°C for 30 minutes, a 1:1 mixture of phenol:chloroform was added and vortexed to homogeneity. After centrifugation, the top layer containing the genomic DNA was removed and mixed with ethanol to precipitate the DNA and pelleted by centrifugation at 16,000 xg for two minutes. The ethanol was decanted and the DNA was resuspended in water and stored at −20°C.

### PCR amplification

Primers for polymerase chain reaction (PCR) were designed for *algT*/*U*, *mucA*, *algO*, and *mucP* (Table 2; Integrated DNA Technologies Inc., Coralville, IA). Each set of primers was designed to fall about 100 bp up and downstream of the gene so that subsequent DNA sequencing would not cut off the beginning and end of each gene.

The *mucA* gene of each of the *sap* mutants as well as the gene identified in the complementation assay as containing a potential mutation were PCR amplified for sequencing. Two and a half micoliters of genomic DNA was mixed with 1 μl of each primer (Table 2), 5 μl of bufferxII, 0.5 μl of HiFi Taq polymerase (Invitrogen, Carlsbad, CA) and the volume was made up to 50 μl with water. The mixture was amplified in the thermocycler with the following program: 95°C for 5 min; 95°C 30 sec, 61°C for 30 sec, 72°C for a time dependent on the amplicon size (1 min/kb) repeated for 30 cycles; 72°C for 10 min; hold at 4°C for further analysis. Products were run on a 2% agarose gel to verify amplification. PCR products were then cleaned according to a kit and standard protocol (Promega, Madison, WI).

### DNA sequencing

Samples were sent to GeneWiz Inc. (Plainsfield, NJ) for sequencing. The *mucA* gene and the complementing genes of each *sap* mutants were sequenced. Samples were prepared according to the company’s requirements. Each tube sent out was premixed with 5 μl of 5 μM primer (forward and reverse separately) and 40 ng of the PCR product.

### Sequence analysis

Sequences were aligned using NCBI Blast, ClustalΩ, T-Coffee, LaserGene software, and Boxshade against the *P. aeruginosa* 2192 wildtype sequence (43, 44) to check for mutations (45-48).

## RESULTS

### Isolation of *sap* mutants

The clinical isolate *P. aeruginosa* 2192 was grown for two weeks at 25°C without aeration. A total of 1058 colonies were analyzed during that period. Of these, 127 (12%) were nonmucoid and considered *sap* mutants. These were then plated on PIA media and allowed to grow for 48 hours to verify stability. Only 41 (3.8%) retained the Alg-phenotype. Of these, 39 *sap* mutants produce alginate when the cells are at high density on a plate and they remain completely nonmucoid when they are single colonies. Two, *sap*8 and *sap*20, remain completely nonmucoid indefinitely, whether in a dense community or single colony.

### Complementation assays

Each of the *sap* mutants was complemented with genes previously identified as common second-site mutations (32, 37). Twenty-eight *sap* mutants (68%) were complemented by pCD100 containing the whole *algT*/*U* operon. These were also complemented by pJG293 containing *algT*/*U* alone (Table 1).

The mucoid phenotype was restored in 11 *sap* mutants (17%) when complemented by *mucP.* Similarly, *algO* successfully complemented four (10%) of the mutants that were also complemented by *mucP*.

Two *sap* mutants, *sap*8 and *sap*20, were not complemented by any of the previously identified genes. When pCD100 (37) was introduced, the two mutants failed to grow. In the presence of pJG293 (37), the strains grew and were nonmucoid.

### Sequencing *mucA*

*P. aeruginosa* 2192 contains a mutation in the anti-sigma factor *mucA* which results in constitutive alginate production. To determine the exact mutation, the *mucA* of *P. aeruginosa* 2192 was aligned with PAO1, the common reference strain which has no mutation, and PDO300, the laboratory-generated strain containing a *mucA22* allele. *P. aeruginosa* 2192 was seen to have A343G resulting in a silent mutation, and G539T leading to a stop codon at the 180^th^ position of the protein (Figure 4). The *mucA* genes of 10 of the *sap* mutants were aligned to *P. aeruginosa* 2192 and shown to possess the original mutations (Figure 4).

**Figure 4:**
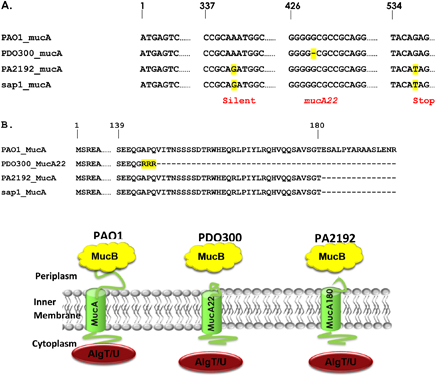
The alignment of MucA. **A.** *mucA* DNA alignment between PAO1, PDO300 (*mucA22*), PA2192, and a sap mutants displaying the differences in *mucA* mutations. Ten *sap* mutants all had the same sequence. PAO1 contains no mutations and is presented as a reference. **B.** Alignment of the respective MucA proteins. **C.** Model of the differences in the MucA protein. MucA22 and MucA180 both have a truncated C-terminus in the periplasm. As a result, MucB cannot bind and the protein loses stability, releasing AlgT/U and bestowing the mucoid phenotype.

### Analysis of *algT*/*U, algO, mucP* in *P. aeruginosa* 2192 and *sap* mutants

#### algT/U

Alignment has shown that the *algT*/*U* ORF and promoter sequence is conserved between PAO1 and *P. aeruginosa* 2192 (data not shown). The *sap* mutants are also mutation free.

#### algO

The *P. aeruginosa* 2192 *algO* sequence shows nine SNPs which all result in silent mutations. The *sap* mutants are yet to be sequenced.

#### mucP

mucP of *P. aeruginosa* 2192 has two SNPs relative to PAO1; one relays a silent mutation, and the other a change of an alanine to a valine in the 313. This change is to an amino acid of similar functional group, and thus it is hypothesized that the function is conserved. The *sap* mutants have not been sequenced as of yet.

## DISCUSSION

Laboratory generated strains have been immensely useful in scientific research and have driven the depth of our knowledge to where it is today. However, laboratory strains, such as PDO300, will always fall short of perfectly mimicking the real-world pathogenesis of clinical isolates (39). This project has certainly confirmed the importance of utilizing a strain isolated directly from the lungs of a patient who passed away from the infection. It has also made the research eminently personal. The present study was designed to confirm the conclusions about alginate regulation drawn from studies using PDO300, as well as investigate the novelty and peculiarity of *P. aeruginosa* 2192.

### *P. aeruginosa* 2912 shows a hyperstable mucoid phenotype

Non-mucoid variants of *P. aeruginosa* 2192 were isolated in the same way as previous studies that utilized PDO300 as the parent strain (37). Studies using PDO300 saw a 90% reversion to *sap* mutants after just 48 hours of culturing at 25°C without aeration (37). In contrast, PA2192 took two weeks under the same conditions to yield even a 3.8% reversion that could be utilized in further analyses. This extended time needed to isolate non-mucoid variants is unique to *P. aeruginosa* 2192 when compared with another clinical strain as well. One study obtained mutants from FRD1, a CF isolate, in 24 hours under the same conditions (32). When compared with the PDO300 and FRD1 studies, *P. aeruginosa* 2192 has a hyperstable mucoid phenotype since it took seven and fourteen times longer before any non-mucoid colony was isolated. It remains to be seen what the contributing factors to this hyperstability are, including the chemical makeup of *P. aeruginosa* 2192 alginate when compared with PDO300. It is interesting to speculate that the hyperstability is directly related to the clinical virulence of *P. aeruginosa* 2192.

### *P. aeruginosa* 2192 possesses a novel *mucA* mutation

Sequence analysis of *mucA* shows that PA2192 does not have the common *mucA22* allele (22) (Figure 4). Instead, it has a previously undocumented *mucA* mutation, which has been named the *mucA180* allele. Alginate production is frequently accomplished in clinical isolates by mutating the *mucA* anti-sigma factor (34). The most common mutation is the *mucA22* allele possessed by ~85% of mucoid *P. aeruginosa* strains (34, 35). It was in this light that PDO300 was constructed from PAO1 with the *mucA22* allele to imitate clinical, mucoid strains (22). The majority of *mucA* mutations are toward the 3’ end of the sequence (24, 49, 50). This results in an altered or truncated C-terminus of MucA that is the end that protrudes from the inner membrane into the periplasm (Figure 4) and interacts with MucB; as a result, MucB binding is reduced or inhibited altogether (50-52). Without MucB binding, MucA is destabilized, and the AlgT/U sigma factor is released resulting in alginate production (50).

It is hypothesized that MucA180 also has a reduced interaction with MucB due to protein truncation. Further experimentation with a yeast two-hybrid system is required to demonstrate this. It could be accomplished by designing a MucA180 bait protein and a MucB fish protein. When compared to wildtype MucA, the MucA180 two-hybrid system should show reduced transcription of the reporter gene.

### *P. aeruginosa* 2192 *algT*/*U* shows no mutation

The *mucA* sequence alignment of the non-mucoid variants revealed that they maintain the *mucA180* allele. Thus, the loss of alginate production is not the result of a true reversion that has restored the function of MucA, but is due to a second-site mutation, as was hypothesized.

Complementation assays have shown that a majority (68%) of the second-site mutations may be mapped to *algT*/*U*. Interestingly, no mutation was detected in the *algT*/*U* open reading frame (ORF) or in the promoter region upstream of the ORF.

This suggests that there may be a novel mechanism of alginate regulation in *P. aeruginosa* 2192 that is bypassed when AlgT/U is overexpressed. Further studies will investigate this possibility. This does not rule out the possibility that this set of revertants may harbor an entirely novel mutation.

### *mucP* and *algO* are involved in *P. aeruginosa* 2192 alginate regulation

This study also found that *mucP* was able to restore the Alg^+^ phenotype of 17% of the non-mucoid variants, indicating that second-site mutations in *mucP* were the second most common mode of alginate repression in *P. aeruginosa* 2192. The third most common second-site mutation in alginate suppression is in *algO* (10%). Moreover, all the non-mucoid variants that were complemented by *algO* were also complemented by *mucP*, suggesting that *algO* mutations can be bypassed in *P. aeruginosa* 2192 by increasing the *mucP* copy number as previously demonstrated (Delgado *et al.*, 2018, Submitted). The exact function of AlgO has not been elaborated as yet but presumed to be a periplasmic protease (37). The *mucP* and *algO* genes in the *sap* mutants are being sequenced to confirm the mutations and find the exact region of the proteins that is mutated.

### Uncomplemented non-mucoid variants possess novel second-site mutations

Two of the non-mucoid variants of *P. aeruginosa* 2192 (*sap2192-8* and *sap2192-20*; Table 1) were not complemented by any of the genes previously identified as liable to second-site mutations. Interestingly, these two also remain completely nonmucoid indefinitely while the others begin producing alginate after 48 hours. This indicates that these possess mutations in one or more novel genes that previously have not been identified as involved in alginate regulation. The *mucA* of these mutants did not possess true reversions and restoration of function. The mutants will be complemented with a previously constructed *P. aeruginosa* cosmid library (37) to identify any novel mutations elsewhere on the chromosome. These two mutants prove fatal when the *algT*/*U* operon was introduced on pCD100. It is hypothesized that either these two do not take up the plasmid for some reason, and thus are killed on the selective media, or the novel mutation, which is yet to be identified, will be able to explain this highly unusual phenomenon.

## Conclusion

Care for CF patients over the last thirty years has dramatically improved. Life expectancy has risen from 18 years in the 1980’s to nearly 50 today (9). *P. aeruginosa*, a ubiquitous bacterium, is devastatingly efficient as a CF pathogen (53). The most indicative factor pointing towards poor patient outcome is the production of alginate by the colonizing strain (54). As of yet, there is no effective anti-alginate therapy. Understanding all variables of *P. aeruginosa* alginate regulation and synthesis is inseparable from combating this deadly pathogen. The present study sought to contribute to this end by investigating regulatory genes involved in the mucoid to non-mucoid reversion of the clinical strain *P. aeruginosa* 2192

## Acknowledgements

We thank members of the Mathee lab for their valuable insights. This research was supported by NIH-National Institute of Allergy and Infectious Diseases (NIAID) 1R15Al111210 (to KM and HK), and NIH-National Institute of General Medical Sciences (NIGMS) T34 GM08368 (to LF). The funders had no role in study design, data collection and analysis, decision to publish, or preparation of the manuscript.

## Ethical Statement

Not applicable.

